# Spider: a flexible and unified framework for simulating spatial transcriptomics data

**DOI:** 10.1101/2023.05.21.541605

**Authors:** Jiyuan Yang, Yang Qu, Nana Wei, Congcong Hu, Hua-Jun Wu, Xiaoqi Zheng

**Affiliations:** Department of Mathematics, Shanghai Normal University, Shanghai, China; Center for Single-Cell Omics, School of Public Health, Shanghai Jiao Tong University School of Medicine, Shanghai, China; Center for Precision Medicine Multi-Omics Research, School of Basic Medical Sciences, Peking University Health Science Center and Peking University Cancer Hospital and Institute, Beijing, China

## Abstract

Spatial transcriptomics technology provides a valuable view for studying cellular heterogeneity due to its ability to simultaneously acquire gene expression profile and cell location information. However, benchmarking these rapidly accumulating spatial transcriptomics analysis tools is challenging owing to the limited diversity and accuracy of “gold standard” data sets annotated by pathologists. To address this issue, we proposed Spider, a flexible and unified simulator for spatial transcriptomics data guided by cell type proportion and transition matrix of adjacent cell types. Taking advantage of a heuristic batched simulated annealing algorithm (BSA) in assigning simulated cell type labels, Spider can generate spatial transcriptomics data for one million cells in just five minutes. Furthermore, Spider can generate various types of spatial transcriptomics data, including immune hot/cold tumor samples by specifying different immune cell proportions and transition matrices and layered tissue samples via an interactive interface. In addition, Spider is also a unified framework for ST data simulation in which we have implemented diverse simulators proposed by other researchers as special cases. We have systematically evaluated the performance of Spider and competing tools, and demonstrated Spider’s remarkable power to capture the spatial pattern of the reference dataset. Spider is available at https://github.com/YANG-ERA/Artist.

## Introduction

Spatial transcriptomics (ST) has revolutionized current investigations of cellular heterogeneity and interplays between gene expression and cellular environment (or position-specificity of gene expression within complex tissues) [1–4]. Unlike single-cell RNA-seq (scRNA-seq) which loses structural information during cell dissociation [5], ST enables quantitatively measuring the expression of individual RNA molecules through spatially indexed barcode and next-generation sequencing techniques [6]. Depending on the experimental assay employed, there are mainly two platforms for performing ST analysis [3]. The first is *in situ* techniques including fluorescence *in situ* hybridization (i.e., seqFISH [7], MERFISH [8], osmFISH [9]), and *in situ* sequencing (STARmap [10] and FISSEQ [11]). Although capable of profiling gene expression at single-cell or even subcellular resolution, this type of method is restricted to preselected encoded probes and unable to achieve whole-transcriptome levels [12]. The second is *in situ* capturing techniques, including ST, 10X Genomics Visium [13], Slide-seq [14], Slide-seqV2 [15], and Stereo-seq [16]. Among them, 10X Visium is more stable, low cost, and has a complete commercial platform, thus becoming the mainstream in spatial transcriptomics research. However, one important limitation of the current 10X Visium technique is lack of single-cell resolution, i.e., a spatial spot with diameter of 55 microns covers approximately 1 to 10 cells.

Based on ST data generated from the above techniques, researchers have proposed over twenty downstream analysis tools [17] to explore novel biological discoveries in plants [18], animals [19], and microbes [20] during the last few years. These tools were developed for a variety of analytic tasks, including denoising [21, 22], identifying spatial variable genes [23–27], detecting spatial domains [28–34], inferencing cell type composition (for 10X Visium and Slide-seq) [35–43], predicting the spatial distribution of RNA transcripts [41, 44-49], identifying cell-cell/gene-gene interactions [50–54], and imputing the missing spatial gene expressions [41, 44, 46, 48, 55, 56]. These tools assist researchers to study the spatial patterns of gene expression in diverse biological settings.

However, the advancement of spatial transcriptomic sequencing technology and analytical methods necessitates a re-evaluation and benchmarking of these tools using gold standard or artificially synthetic datasets. Gold standard data refer to ST data with known spatial domain or cell type annotation by pathologists. Despite the accumulation of ST data in recent years, such as the mouse hypothalamus data [57] by MERFISH and human dorsolateral prefrontal cortex data [58] by 10X Visium, researchers are still facing the following limitations: i. Cell type annotation by pathologists is not guaranteed to be the ground truth, because pathologists cannot effectively combine cellular phenotypes (e.g., cell size, shape, and nuclear density) with molecular profiles [4]. ii. The number of benchmark datasets and their volumes are insufficient to validate the computational efficiency of different methods; iii. Available benchmark datasets exhibit a preference for tissue samples that possess a distinct layered structure, while disregarding tumor samples showing spread or mutually exclusive patterns of infiltrated immune cells [59]. These tumor samples are challenging to manually annotate but may hold greater significance in clinical contexts. Thus, a feasible way is to utilize simulated data instead to overcome these limitations and boost the ST method development and evaluation [60, 61].

We summarized the following four critical criteria for an ideal ST data simulator: i. Flexible spatial patterns. The simulator can generate synthetic data with varying spatial patterns; ii. Interactive layer customization. The simulator provides an interface for the user to tailor the layer structure and to dynamically manipulate the shape and dimensions of the spatial domain; iii. Easy to implement. The simulator should be open source and provide user-friendly API for redevelopment; iv. Computational efficiency. The simulator should not require expensive hardware support or excessive computational time. However, to the best of our knowledge, none of the existing simulators embrace the above four criteria simultaneously. They either oversimplify the ST data by disregarding several crucial features (such as the tendency for adjacent spots to exhibit greater similarity in expression levels [58]) or lack proper documentation and package for reproducibility, which hinders the broad evaluation of ST data analysis tools.

Here, we developed Spatial Transcriptomic Data Simulator (Spider), a Python-based framework for reproducibly simulating spatial transcriptomic data (Fig. 1). A key feature of Spider is to use prior distribution and transition matrix of cell types to characterize the spatial pattern of simulated data. In particular, Spider can generate ST data with a hierarchical structure in an interactive manner. Moreover, Spider provides an implementation interface for most existing simulation methods and a collection of available gold-standard datasets facilitating researchers in promptly characterizing, benchmarking, and evaluating new analytical methods. Compared with existing simulation methods, Spider shows superior performance in terms of both consistencies to reference data and computational efficiency. Additionally, Spider exhibits the capacity to delineate immune hot/cold tumor samples and stratified tissue samples that encompass diverse cell type compositions and spatial patterns.

**Fig 1.**
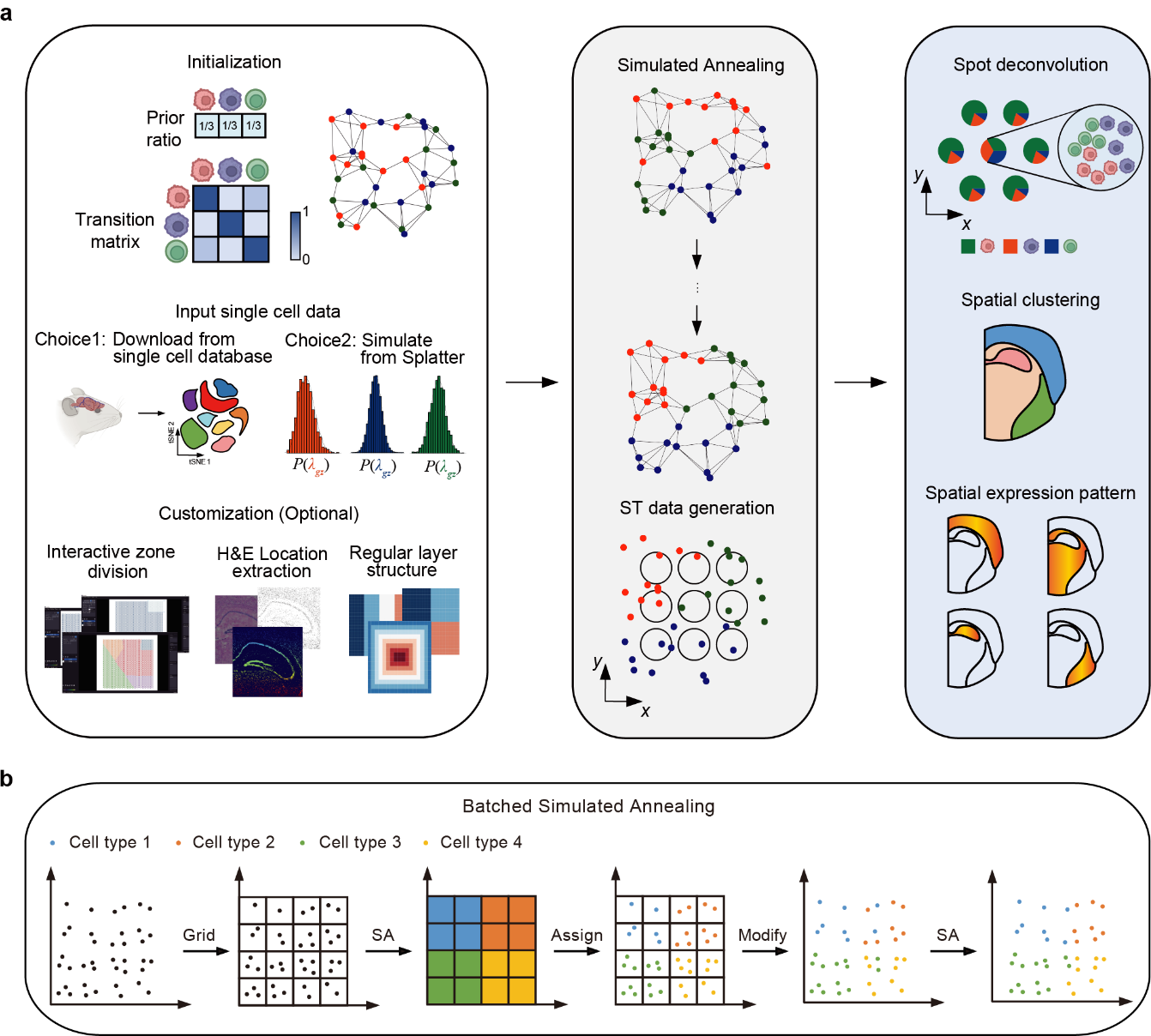
| Framework of Spider. a. Spider is a software framework for simulating ST data. Input parameters are indicated with solid border white background. Spider begins by inputting the prior ratio, transition matrix, and number of cells and constructing a spatial neighbor network. Cell expression profiles are generated by real data or simulated from splatter. Adjusting the cell type label using the simulated annealing algorithm enforces to approach the targeted transition matrix. In the final step, spot-level ST data is generated by a setting regular spot array like 10X Visium. To obtain the gene expression profiles at each spot, Spider sums the gene expression profiles of all cells in one spot. Evaluating the methods of downstream analysis is indicated with solid border blue background. In addition, the bottom left corner displays optional applications of Spider, such as interactive zone division and cell location information extraction from H&E image, and offers a variety of standard layer structures. **b.** Batched simulated annealing algorithm in Spider. Similar to the input of the Spider, after determining the spatial position matrix of the cells and the network relationship, we grid the simulated cell space regions and randomly assign the cell type labels to each grid based on the prior cell type proportion; then, Spider regards this grid as the representative unit of the cells in this area. We treat all representative units as a batch and perform the simulated annealing algorithm; According to the label of the representative unit, Spider retraces and assigns the same label to all cells in the corresponding region; Finally, Spider performs the simulated annealing algorithm at the cell-level scale until the cell type arrangement satisfies the user-specified transition matrix.

## Results

### Overview of Spider

Here, we developed Spider, a universal simulation framework for ST data with various cellular spatial patterns (e.g., tumor immune microenvironments, tissue types, and developmental stages). Briefly, Spider consists of three steps: (1) model parameter initialization; (2) cell type assignment and gene expression allocation; (3) Aggregation to spot level (Fig. 1a).

In the first step, users need to specify the number of cell locations, the number and proportion of cell types, an optional transition matrix between cell types and matched scRNA-seq data. Not only can the transition matrix be replaced by a specified spatial pattern, including addictive, exclusive, and striped (see Supplementary Material for details), scRNA-seq can also be generated from Splatter [62] or scDesign2 [63]. Then, Spider generates locations of cells on a plate randomly or in a uniform grid-like pattern. We next construct a neighborhood graph of cell locations according to their pairwise distances on the plate. Spider now supports various neighborhood metrics, such as k-nearest neighbors [64] or neighbors identified by Delaunay triangulation [65, 66], a more accurate representation of cell neighborhoods in real settings.

In the second step, Spider aims to assign the cell type labels to approximate the user-specified spatial pattern. We formulated it as a binary optimization problem with integer constraint (see Method for detail). However, the cell type assignment problem is difficult to solve analytically, we thus proposed a heuristic batched simulated annealing (BSA) algorithm (Fig. 1b), motivated by the convolutional neural network (CNN) [67] in deep learning, to adjust cell type labels in parallel. Briefly, Spider partitions the cell plate into several batches and employs the simulated annealing algorithm to converge the transition matrix on the batch level to the user-specified transition matrix. As the batch size decreases, the global transition matrix gradually converges to the target, leading to a significant enhancement in convergence speed (Fig. 1b).

In the last step, Spider generates the synthetic ST data at the spot level by aggregating gene expressions of all cells within each spot. Users can specify the shape (circle or rectangle) and size of spots to generate a regular array of spatial spots, then the expression profiles of all captured cells within each spot are summed up to obtain the spot-level expression profiles.

In addition, Spider also supports optional interactive operations in ST data simulation, including domain segmentation, extraction of cell coordinates from hematoxylin and eosin (H&E) images, and domain structure customization. For interactive domain segmentation, Spider can manually delineate the spatial domain of simulated cells with the help of *Napari*, a multi-dimensional image viewer in the Squidpy [66] package. Spider can also take H&E staining images as input to generate cell coordinates using an anatomical segmentation algorithm Cellpose [68], which is specifically designed for nuclei segmentation. At last, Spider provides several customized regular spatial domain structures for users, such as blocked, striped, and gyrate (Additional file 1: Fig. S1).

### Summary of existing ST simulation tools

We systematically summarized the existing simulation algorithms, including Spider, and divided them into three main categories based on the underlying principles (Fig. 2a-c). The first category is random-based simulations, which randomly sample cells from real scRNA-seq data (or simulated data from Splatter [62] or scDesign2 [63]) and assign them to each spot with different locations to generate ST data (Fig. 2a). Methods in this category include stereoscope [40], RCTD [35] and STRIDE [36]. These approaches can generate a set of ST data fast, but they do not take into account the spatial dependence of cell types and gene expression associations among adjacent cell locations. The second category is aggregation-based methods, which first generate spot-level ST data by aggregating a single-cell resolution spatial transcriptomic data, where gene expression of each spot is generated by summing up all cells within that spot (Fig. 2b). Methods in this category include STdeconvolve [37], SpatialPCA [69] and BASS [29]. This scheme preserves the spatial pattern of the original data but relies on ST data with single-cell resolution, which is not flexible enough to reflect diverse spatial patterns or control the number of cells for benchmarking. The third strategy is model-based simulations, which are based on different statistical models of data generation (Fig. 2c). Methods in this category include FICT [30] and cell2location [39]. Some methods depend on certain model assumptions, which may be not generalizable to other datasets. It is worth mentioning that none of the aforementioned methods is specifically designed for simulating ST data, but to validate their downstream methods. To enable researchers to utilize these methods within a unified framework, we also implemented all these methods in Spider. Users can easily use the above-mentioned simulators by setting the proper parameters. For example, one can apply RCTD by simply executing the function *spider.rctd(spot_num, celltype_list, min_cell, max_cell, reference)*, in which *spot_num* represents the number of spots simulated, *celltype_list* represents the cell type specified for simulation, *min_cell*/*max_cell* represents the minimum/maximum number of cells contained in the spot, and *reference* represents the reference expression profile.

**Fig. 2.**
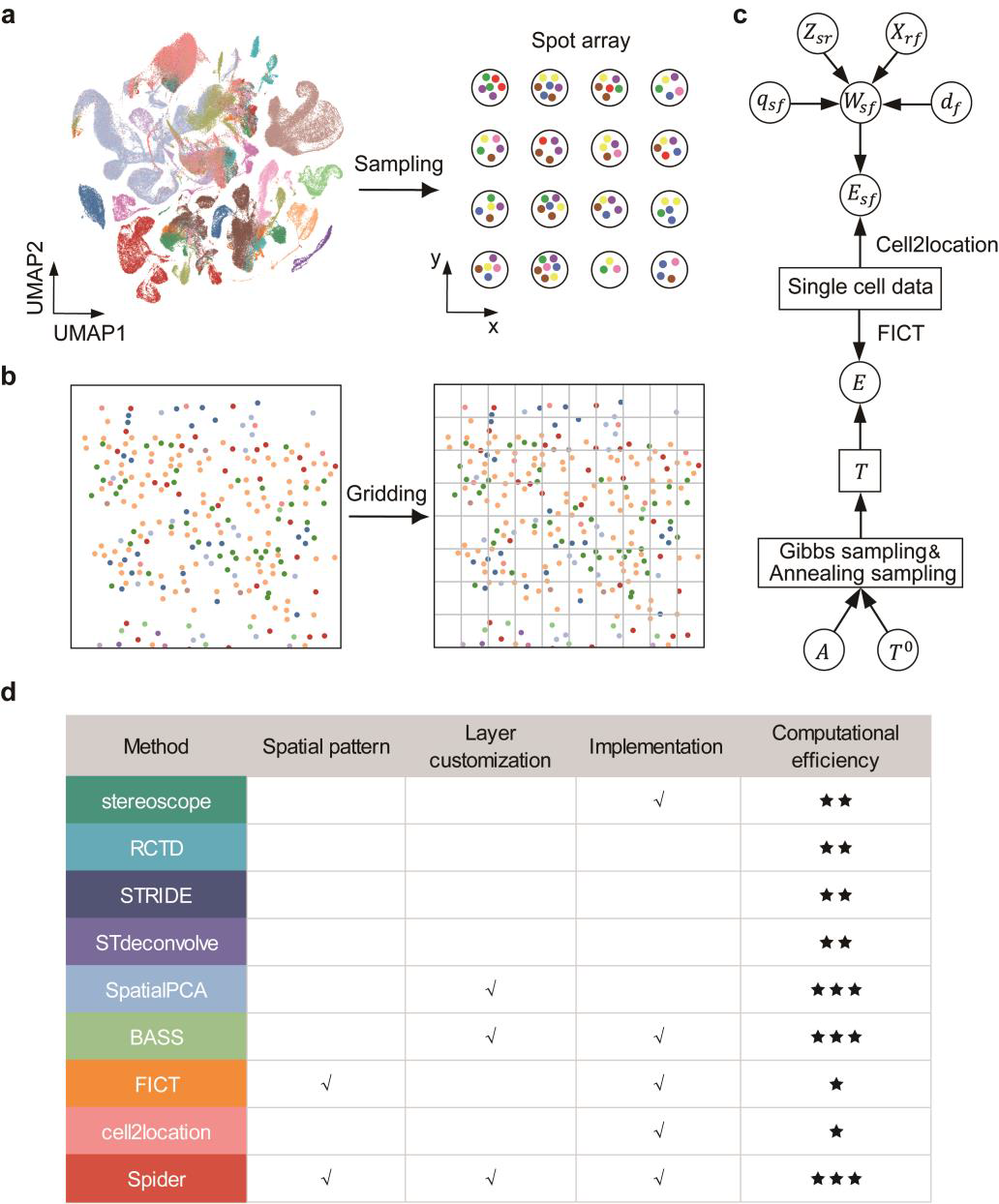
Summary of existing simulation methods. a. Random-based. For each spot, according to the distribution assumption, some cells of different cell types are extracted from the scRNA-seq database and placed into the spot, and the gene expression values of each spot represent the sum of all the cells in that spot. **b.** Aggregation-based. It generates data by gridding the single-cell resolution spatial transcriptomic data, where each grid represents a simulated spot containing multiple cells. **c.** Model-based. Here, we listed the most typical algorithms such as FICT, and Cell2location, and use the probability diagram to show their process. **d.** Summary of nine simulation methods against four key criteria. For the computational efficiency column, the score depends on the running time on the test data with a sample size of 10,000, which means: 3 stars: Running time less than 30 seconds. 2 stars: Running time between 30 and 120 seconds. 1 star: Running time more than 120 seconds.

We next compared nine representative simulators in Fig. 2d listing their ability to fulfill four key criteria. Two model-based simulators, FICT and Spider, preserve spatial patterns through a user-specified transition matrix between cell types, thus satisfying criterion 1. Although STdeconvolve can preserve the spatial distribution of reference data, it is limited by reference ST data with single-cell resolution and cannot flexibly generate diverse data. SpatialPCA and BASS change the proportion of cell types within each domain and the size of the spatial domain, then randomly assign each single cell in each domain to one of the cell types based on a multinomial distribution with parameters set to be the cell-type composition in the domain, thus satisfying criterion 2. Spider can not only set different cell type proportions in each spatial domain as adopted by BASS, but also can interactively select spatial domains, or use anatomical reference atlas pictures as the background, which is convenient for users to generate tissue structures with different layers of cell types, thus satisfying criterion 2. RCTD, STRIDE, STdeconvolve, and SpatialPCA only presented their methodologies as pseudo-codes and no implementation was provided, which violates criterion 3. Finally, we evaluated the computational efficiency of the simulation methods. Model-based methods, such as FICT and cell2location, generate data from statistical models. As a result, the computation times are generally higher than those of random-based methods. FICT utilizes simulated annealing algorithms to optimize the spatial arrangement of cell types [30]. This is achieved by randomly switching pairs of cells, which results in high computational demand when simulating datasets exceeding 10,000 cells. Although Spider also uses simulated annealing, it achieves improved computational efficiency even for tens of thousands of cells by taking advantage of the BSA algorithm.

### Performance comparison of ST simulation tools

We next evaluated different ST simulation tools based on their agreements with real spatial transcriptomic data. To achieve it, we downloaded single-cell resolution mouse visual cortex data, which captures 1549 cells corresponding to 15 cell types obtained through clustering with careful marker gene examination [70]. We removed the ambiguous cell types and retained six cell types with sufficient number of cells as cell-level reference data, including astrocytes (141 cells), endothelial (150 cells), excitatory L2/3 (258 cells), excitatory L4 (198 cells), excitatory L5 (94 cells), excitatory L6 (287 cells) (Supp file 1: Fig. S1). Then the spot-level data is generated by the aggregation-based method as ground truth. Specifically, we gridded the cell level (single-cell resolution) data with squared windows of 500 x 500 pixels, resulting in 347 spots with each spot consisting of 1–10 cells (Additional file 1: Fig. S2). Each simulated spot contains only one cell randomly sampled from the reference dataset to reasonably simulate the cell-level data using random-based methods. Based on the reference dataset, we estimated the prior cell type proportion and transition matrix for Spider.

We evaluated the performance of different methods in terms of consistency with real data from the following two aspects: cellular spatial patterns and gene expression spatial patterns. The reference dataset shows a cluster of excitatory neuron cells between layers 2 and 3 (termed Excitatory L2/3). These excitatory neurons are primarily surrounded by endothelial cells (Fig. 3a, left panel). This structure is exclusively observed in the dataset generated by Spider (Fig. 3a, the second panel). We next evaluated the spatial similarity of the simulated data produced by each method in terms of the consistency of the transition matrix (TM), centrality scores matrix (CSM), and neighborhood enrichment matrix (NEM) (see Methods, Fig. 3b). The errors of each method are calculated as the distance between real and estimated matrices. To obtain more robust results, we repeated the simulations of each method 10 times. Among the methods tested, Spider exhibited the lowest error between the synthetic and real data, indicating its ability to capture the cellular spatial pattern of the reference dataset.

**Fig 3.**
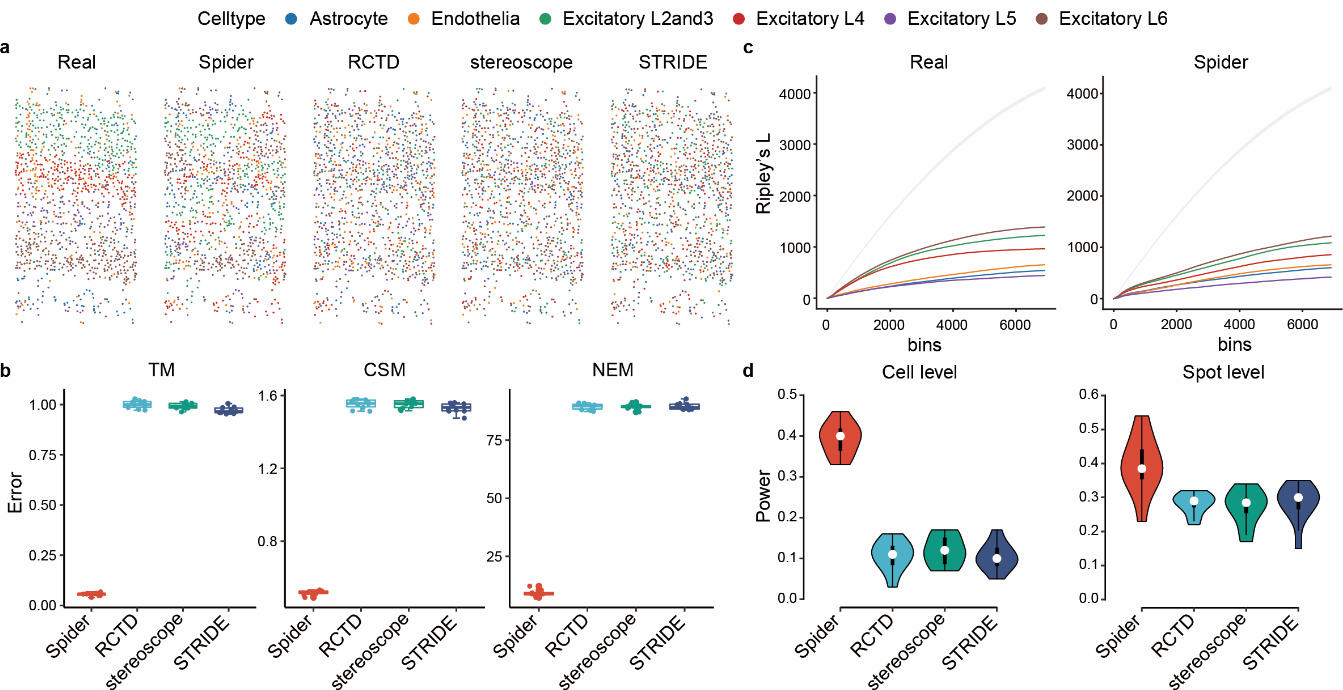
| Comparison of ST simulation tools. a. The spatial distribution of the STARmap dataset and simulated results (Spider, RCTD, stereoscope, STRIDE). Excitatory L2/3, 4, 5, and 6: excitatory neurons in layer 2/3, 4, 5, and 6. **b.** The boxplot of the error about spatial pattern statistics (transition matrix, centrality scores matrix, neighborhood enrichment matrix) compared with real data and simulated data generated by other methods. **c.** The Ripley’s *L* curve shows the degree of deviation of cell-type-specific density distribution from the null hypothesis of random point pattern, as shown in both the reference dataset and the simulated data by Spider. The gray-shaded line represents the expected L-function of a random point pattern. **d.** The violin plot of benchmark SVGs power for synthetic data generated by four simulation methods.

To further assess the similarity of the distribution of all cell types, we plotted the Ripley’s *L* curve for each method. This allowed us to detect whether the six cell types showed a global pattern, e.g., random, clustered, and dispersed [71] (Fig. 3c). It is clear that all cell type clusters simulated by Spider are below the baseline, indicating a dispersed pattern across the area. The same phenomenon is observed in the reference dataset, indicating that Spider accurately captures the spatial distribution characteristics of cell type density in the reference dataset. Although the cell type clusters simulated by RCTD, Stereoscope, and STRIDE are also below the baseline, most of their cell types show a significantly different trend in Ripley’s *L* value compared to the reference dataset across increasing distances (Additional file 1: Fig. S3). Next, we compared different methods for recovering gene spatial patterns (Fig. 3d). Based on the reference data, we selected the top 100 genes with the largest Moran’s *I* spatial autocorrelation statistics as benchmark spatially variable genes (SVGs) [66, 72]. We then computed the power value, which represents the ratio of overlap between the SVGs identified in each simulated data and those identified in the real data, to evaluate whether the simulated data could capture the spatial patterns of genes. In both cell-level and spot-level data, Spider shows the highest power value, which is almost three times that of RCTD, stereoscope and STRIDE, and preserved 40% of the spatial expression differences of the benchmark SVGs (Fig. 3d). Overall, Spider is effective and robust in preserving the spatial pattern of the reference dataset.

### Application of Spider in tumor and tissue simulation

In this section, we showed the application of Spider in generating layered tissue samples and tumor samples with different immune microenvironments (TIME) [73]. TIME has been an important research topic in cancer immunology [74]. Previous studies have shown that different tumor cells secrete various cytokines and chemokines, which can create a local immune microenvironment by recruiting different types and amounts of immune cells [75–78]. It has been reported that the spatial organization of immune phenotypes can be classified into three categories: cold, mixed and compartmentalized [79]. Immune cold tumors show few immune cells infiltration, mainly macrophages (Fig. 4a). Mixed tumors exhibit a high amount of immune cells, and tumor and immune cells are mixed (Fig. 4b). In compartmentalized tumors, immune cells are present in high proportion but are spatially segregated from tumor cells (Fig. 4c). For example, neutrophils tend to be enriched near the border, while B cells form secondary lymphoid structures further away.

**Fig 4.**
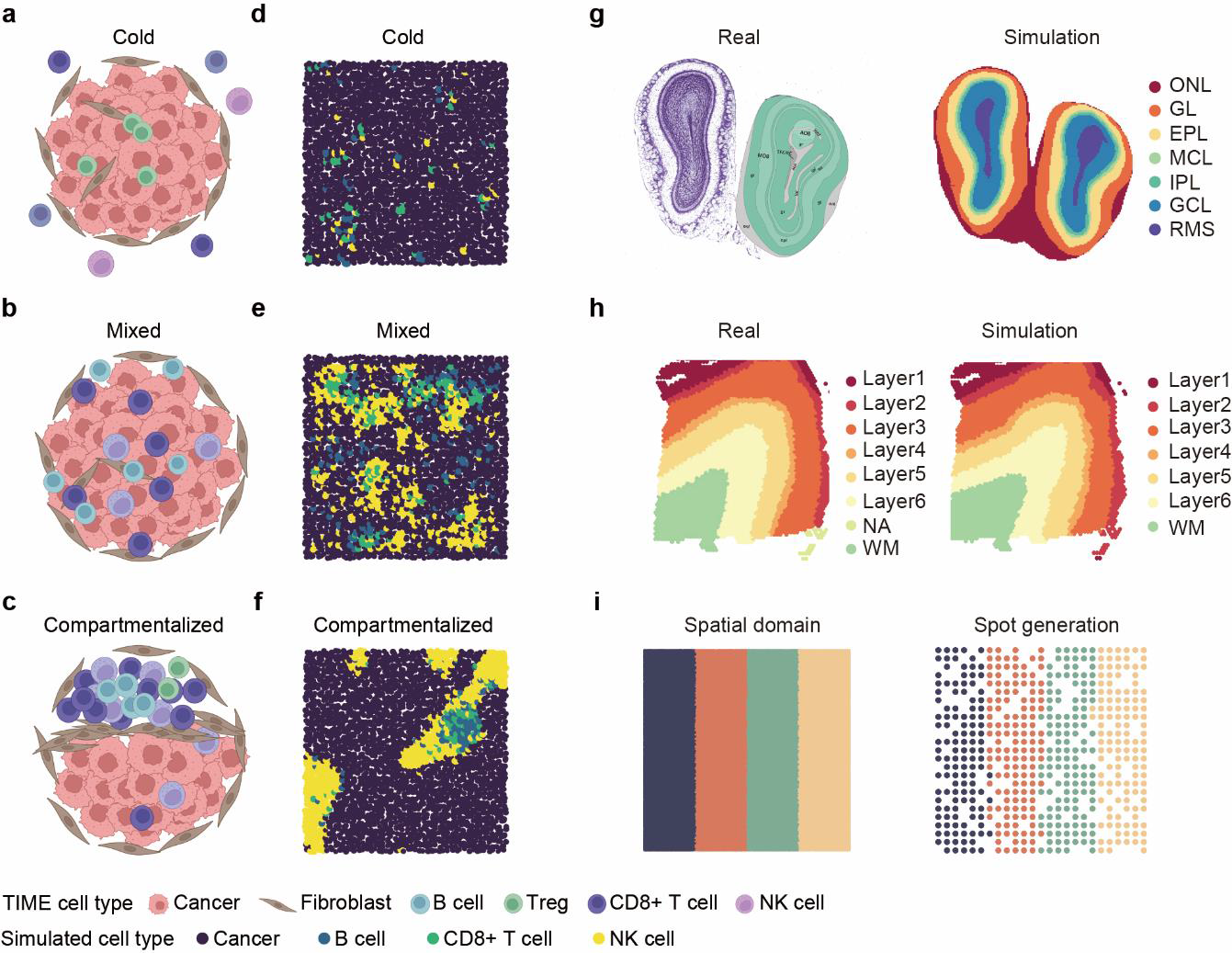
| Result of ST data simulation. a-c. Three archetypes of the tumor-immune composition of ST data (cold, mixed, and compartmentalized). **d-f.** Corresponding simulation result of three archetypes of tumor-immune composition. **g.** Real spatial structure of mouse olfactory bulb (left) and the simulated result of mouse olfactory bulb using interactive method (right). **h.** Real spatial structure of DLPFC (left) and the corresponding simulated result using interactive method (right). **i.** Result of simulation by generating spatial structure of simple patterns.

We applied Spider to simulate ST data with three types of TIME by setting proper cell type proportion and transition matrix (Fig. 4d-f, see Supplementary for detail). Mixed tumors simulated by Spider reveal that a majority of immune cells infiltrate the tumor cells (Fig. 4b), which are dispersed throughout various regions of the tumor. Additionally, certain immune cells exhibit characteristics of aggregation, closely resembling the observed TIME. Similar trends are observed in cold and compartmentalized simulated tumors. This result indicates that Spider can effectively simulate the diverse tumor-immune spatial landscapes by adjusting the cellular transition matrix and cell type proportions.

In addition, Spider provides tissue simulations with distinct layered structures in an interactive manner. In detail, we can first generate a regular lattice according to the given interval between spots by Squidpy package. Users can manually divide similar layer structures on the lattice according to the biological structure of the target tissue, and generate layer labels. Based on generated label and location information, Spider generates corresponding tissue ST data. As an example, we generated simulation data based on existing tissue information of the mouse olfactory bulb [16, 80] and DLPFC (Fig. 4g-h), and the obvious layer structure is well preserved. Additionally, we also generated ST data with regular patterns (Additional file 1: Fig. S4). In conclusion, Spider can generate versatile and diverse ST data, which presents significant value for researchers to evaluate novel algorithms.

### Evaluation of downstream spatial transcriptomic methods

To further demonstrate the effectiveness of simulated data, we conducted downstream analyses on both real and simulated ST data generated from various methods, including Spider, FICT [30], RCTD [35], Stereoscope [40], and STRIDE [36]. Our focus is primarily on two types of downstream analysis tasks: spatial domain detection (clustering) using SpaGCN [32], SC-MEB [34], BASS [29], and BayesSpace [28], and spot deconvolution using STRIDE [36], SpatialDWLS [81], RCTD [35], Tangram [45] and Stereoscope [40]. We reason that if a simulation method can effectively capture the spatial pattern of real data, the simulated data should show a similar pattern to that of real data. To achieve this, we used the STARmap data as a reference and generated spot-level data by gridding and aggregating procedures illustrated in the previous section. Then, we generated simulated data using different methods according to the STARmap data (see Methods for details). Each simulation is repeated 10 times to minimize randomness.

We began by comparing different deconvolution analysis results on the obtained simulated data. To quantify the accuracy of deconvolution, we measured the average correlation between the estimated cell-type compositions and the ground truth across all spots. Compared to the other simulation methods, the simulated data produced by Spider consistently yield deconvolution results that were comparable to real data, regardless of the deconvolution approaches used (Fig. 5a). In particular, all deconvolution methods perform poorly on data simulated by STRIDE, probably because STRIDE ignores spatial dependence due to its random sampling. Based on our evaluation of the simulated data, RCTD and Stereoscope yield comparable accuracy and outperform other methods, consistent with the conclusions drawn by Li et al [82].

**Fig 5.**
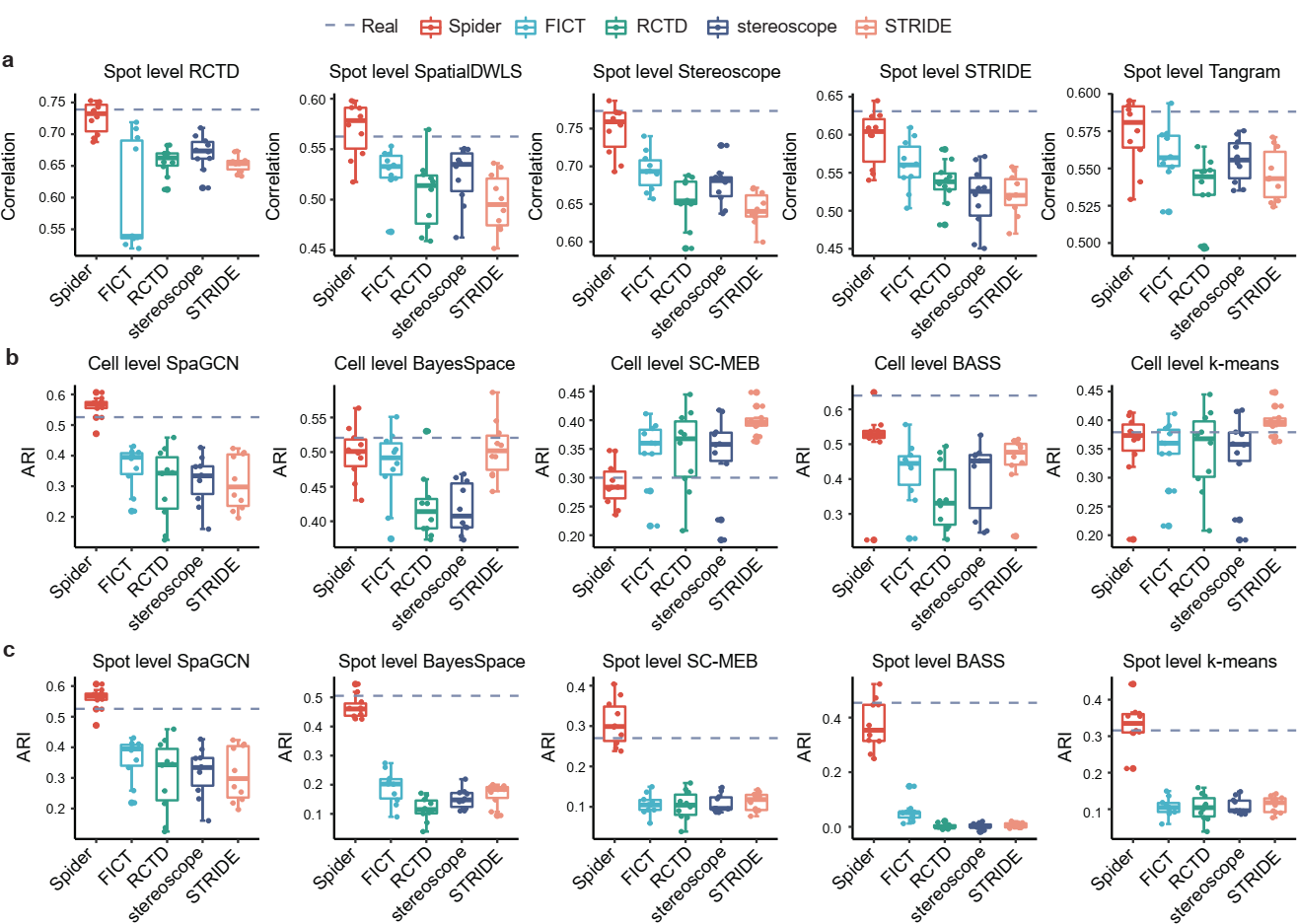
| Benchmarking downstream analysis methods on different simulated data. a. The correlation between the deconvolution results on the data generated by the five simulation methods and the real data. **b**-**c**. The accuracy of different clustering methods on the various simulation data and real data, where each simulation data corresponds to two-level data, including cell (**b**) and spot (**c**). The gray dotted line represents the result of the real data.

Next, we compared different spatial clustering results on the obtained simulated data. We used K-means as a baseline method and measured the adjusted Rand index (ARI) [83] to evaluate the clustering accuracy. The results on the simulated data from our method are more consistent with the real data, in both cell-(Fig. 5b) and spot-levels (Fig. 5c). Especially for spot-level data, our method shows significantly higher agreement with real data than other simulated methods. As expected, due to overlooking the spatial dependence, the average ARI of K-means is lower than other clustering methods that incorporate spatial correlation information. These results demonstrate that Spider can produce simulated data that accurately captures spatial information.

## Discussion and conclusions

In this article, we present Spider, a comprehensive framework for simulating ST data by leveraging spatial neighborhood patterns among cells through a conditional optimization algorithm. By inputting spatial patterns extracted from real data or user-specified transition matrix, Spider assigns cell type labels through the BSA algorithm that improves the computational efficiency of conventional simulated annealing algorithm [84]. Following this process, gene expression profiles for individual cell types can be generated either using real or simulated data by Splatter or scDesign2. Finally, spot-level ST data can be simulated by aggregating cells inside generated spots using 10X spatial encoding techniques [13] or user-specified generation rules.

Different from existing simulation methods, such as RCTD, STRIDE, and STdeconvolve, Spider is driven by the characteristics of ST data and spatial distribution patterns of cell types. These results in more realistic simulated data and the ability to flexibly model various scenarios. In addition, we have implemented all the simulation algorithms mentioned in the paper using Python in Spider to make our framework unified.

We used the STARmap dataset with the single-cell resolution to evaluate the performance of Spider and the other existing simulation methods on cellular spatial patterns and gene expression spatial patterns. Overall, Spider achieves excellent performance and robust results in preserving the spatial patterns in the STARmap dataset. We also conducted downstream analysis, including spatial clustering and spot deconvolution, on the ST data generated by different simulation methods. The results obtained on the Spider simulated data are highly consistent with the reference data. This indicates that Spider preserves the spatial features and different cellular neighborhood patterns during simulation.

Thanks to the capability of Spider to synthesize data by creating a transition matrix between cell types, the simulation of spatial transcriptomic data for tumor immune microenvironment has become achievable. By simulating various scenarios of the tumor immune microenvironment and capturing the dynamic changes in different immune cell components, Spider can provide robust support for researchers developing algorithms aimed at studying complex tumor ST datasets. In addition, Spider has additional customized data generation APIs for special application scenarios, such as tissue layer structure implemented by *Napari* interface. If users need more accurate cell location information, Spider can also extract it from the H&E image. At the same time, we also provided some regular structures for reference. In conclusion, Spider’s flexible framework enables researchers to better simulate ST data in different scenarios, offering enormous potential to explore the spatial expression patterns of genes in biological tissues, revealing the complexity of biological systems, and facilitating the development of downstream methods.

During our article submission, we discovered two recent studies that employ similar methods to simulate ST data. Baker et al. [61] utilized probabilities of cell-type discovery and transition matrices to regulate spatial patterns. However, they did not successfully address the issue of assigning cell-type labels to the cell plate. On the other hand, Zhu et al. [60] developed SRTsim that can simulate ST data by preserving the spatial gene expression patterns, but is limited in terms of generating diverse simulations and is unable to simulate tumor ST data with multiple states due to the absence of reference.

Besides ST data, the emergence of spatial omics technologies has enabled the spatial resolution of proteins, metabolites, and DNAs in unprecedented detail in recent years [59]. In our future work, we will endeavor to incorporate various developed spatial transcriptomic simulation algorithms into the framework of Spider, and expand Spider to the simulation of other spatial omics data (e.g., Spatial Proteomics [85–87], Spatial Metabolomics [88]), providing convenient data support and potential biological insights to bioinformatics researchers.

## Methods

### Cell type assignment

Spider is designed to simulate ST data with specific spatial patterns. It first generates a single-cell resolution ST data, assuming it contains *N* cells belonging to *K* different cell types, and then generates spot-level data via a grid-merge approach, where the expression profile of each spot is rendered in the total composition of the expression profiles of the cells. Spider generates single-cell resolution data as following: given the number of cells *N*, the spatial coordinates of each cell are first generated from the uniform distribution, denote *S* = (*s*_1_, …, *s*_*N*_)^*T*^, *s*_*i*_ ∈ *R*^2^/*R*^3^. Then, assign a cell type to each cell, and obtain expression profile data for each cell sampling from single-cell data or distributions of different cell types according to the cell type labels. Among them, how to assign cell types reasonably is crucial. We achieve the assignment of different spatial patterns by setting the transition probability among cell types.

Given the prior proportion of initial cell types *π* = (*π*_1_, …, *π*_*K*_)^*T*^, where 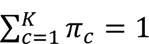, the transition probabilities between cell types *P* ∈ *R*^*K*×*K*^ . The final cell type labels *X* ∈ *R*^*N*×*K*^ should satisfy the following conditions: the transition probability between cell types after assigning cell types is as close as possible to the given transition probability, and the proportion of each cell type agrees with the given proportion. Thus, we have the following optimization objective:

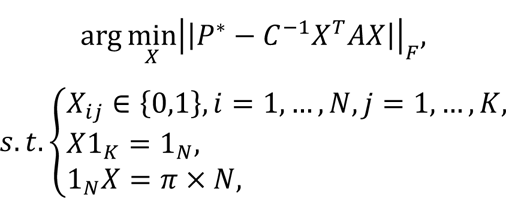

where *A* is the adjacency matrix of the graph built by spatial coordinates *S* of cells (see Spatial graph construction below). *X*^*T*^*AX* is the transition frequency matrix based on the cell type assignment *X*. *C* = *diag*(*X*^*T*^*AX*1_*K*_) is the row-normalized factor matrix. The cell *i* belongs to the cell type *j* if *X*_*ij*_=1.

### Batched Simulated Annealing

However, solving the above constrained binary optimization problem is computationally infeasible, and it is also unrealistic to calculate the optimal solution by grid search when the dimension of the variables is large. When the cell type *K* exceeds 2, the cell type label assignment problem is NP-hard. To address this issue, we developed an efficient batched simulated annealing algorithm (BSA) based on simulated annealing, a heuristic search technique, to find an acceptable solution [84].

BSA performs simulated annealing on cell zones rather than on each cell. The BSA algorithm mainly involves three phases: (1) initialization of the cell zones and cell type labels; (2) implementation of simulated annealing algorithm; (3) subdivision of each cell zone into sub-zones. Repeated (2) and (3) until the given termination condition is satisfied, culminating in the final categorization of cell types for every individual cell. Specifically, the cell plate is regarded as a rectangular area in two-dimensional space. The entire region is initially divided into *l* ∗ *w* grids, with each grid representing a cell zone, where *l* and *w* represent the number of bins divided by length and width, depending on the number of cells. Cell types in each zone are then sampled using a multinomial distribution based on the initial cell type proportions *π*. Subsequently, the simulated annealing is employed to swap cell-type labels across distinct regions, thereby enabling the convergence of the cell-type transition probability matrix to the target transition probability matrix. Next, the cell zone is divided into sub-zones of size ℎ ∗ ℎ (default by 2). We assumed that the initial cell type labels of each sub-zone are consistent with the label of its corresponding zone. The cell type reassignment of sub-zones is performed by simulated annealing. Iteratively subdivide the zones until reaching the cell level. Compared with directly performing simulated annealing on the original resolution data set, the BSA algorithm can greatly improve the convergence speed.

### Spatial graph construction

Spatial patterns are encoded in the construction of spatial networks that reflect relationships between neighboring cells. Each node in the network represents a cell with its specific attributes, and connections between the nodes represent the neighbor relationships among surrounding cells. The construction of various types of networks plays a fundamental role in calculating the subsequent transition matrices, which help to capture the complex spatial interactions between cells [66]. We construct an undirected graph through the Euclidean distance between any two cells using spatial information *S*. The adjacent matrix *A* can be calculated as:

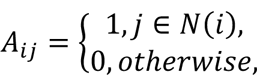

where *N*(*i*) represents the set of neighbors of cell *i*.

### Benchmark metrics

To evaluate the performance of the simulation methods. We used the following three metrics to benchmark these simulation methods.

#### 1. Clustering accuracy

ARI and NMI. We use adjusted Rand index (ARI) [83] and normalized mutual information (NMI) [89] to measure the clustering accuracy of different methods, which are commonly used in evaluating the clustering performance. Given a set of *n* spots and two clustering labels for these spots, the ARI is calculated as:

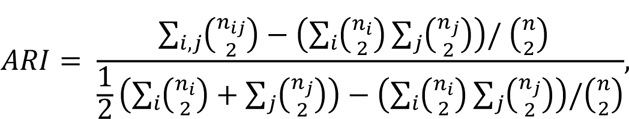

where *n*_*ij*_ is the number of spots overlapped by cluster *i* and cluster *j*. *n*_*i*_ and *n*_*j*_ are the number of spots in cluster *i* and *j*, respectively.

The NMI is calculated as:

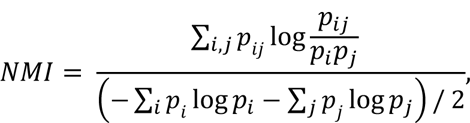

where 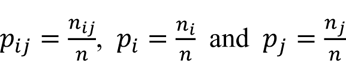.

#### 2. Spatial pattern metrics

We used the three matrices, transition matrix (TM), centralized score matrix (CSM), and neighborhood enrichment matrix (NEM), to measure the spatial pattern [90, 91]. Given a graph *G* = (*V*, *E*) based on an ST data, where *V* and *E* represent all *N* spots and edges between spots, respectively. Let *C*_*k*_ be the group of spots with the same cell type. TM is the row normalized transition frequency matrix.

The CSM is used to describe complex relationships in large networks and can be represent as:

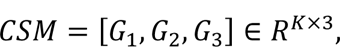

Here *G*_1_ is group degree centrality, measuring the ratio of non-group members that are connected to group members. The formula is described below:

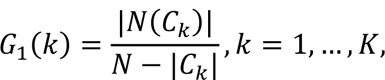

where *N*(_*k*_) denotes the neighbors of all nodes in *C*_*k*_. *G*_2_ is the average clustering coefficient measures how likely the cluster nodes favor to cluster together. The formula is described below.

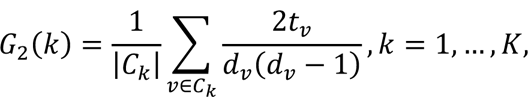

where *t*_*v*_ denotes the number of triangles around node *v* and *d*_*v*_ is the degree of node *v*. *G*_3_ is the group closeness centrality is defined as the normalized inverse sum of distances from the group to all nodes outside the group. The formula is described below:

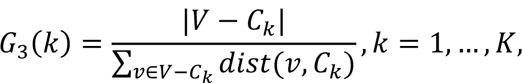

where *Cist*(*v*, _*k*_) = min_*v*′∈_ *Cist*(*v*, *v*^′^) denotes the shortest distance between the group _*k*_ and node *v*.

NEM quantifies the enrichment between each pair of cell types [92]. Let *x*_*ij*_ be the number of connections between _*i*_ and _*j*_. Through preserving the spatial structure and shuffling the cell type labels *m* times, and then recounting the number of connections between each pair of clusters, a z-score can be calculated as:

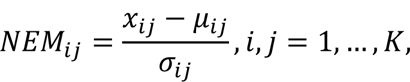

where *μ*_*ij*_ is the expected means and *σ*_*ij*_ is the standard deviations for each pair in the random configuration.

For both simulated and reference data, we can obtain the above 3 matrices. The Frobenius norm of the difference between two matrices is calculated to assess the closeness of the spatial pattern of cell types between the simulated and real data.

## Data availability

All the datasets analyzed are publicly available via respective publications. The STARmap dataset (mouse visual cortex) is available at https://www.starmapresources.com/data; Smart-seq, (mouse primary visual cortex) is available at https://portal.brain-map.org/atlases-and-data/rnaseq/mouse-v1-and-alm-smart-seq.

## Code availability

Spider is a Python package at the following Github repository: https://github.com/YANG-ERA/Artist. All data synthesized by different methods can be found in https://github.com/YANG-ERA/Artist. All the code to reproduce the result of the analysis can be found at the following GitHub repository: https://github.com/YANG-ERA/Artist.

